# Genetic mapping of species differences via “*in vitro* crosses” in mouse embryonic stem cells

**DOI:** 10.1101/148486

**Authors:** Stefano Lazzarano, Marek Kučka, João P. L. Castro, Ronald Naumann, Paloma Medina, Michael N. C. Fletcher, Rebecka Wombacher, Joost Gribnau, Tino Hochepied, Claude Libert, Yingguang Frank Chan

## Abstract

Discovering the genetic changes underlying species differences is a central goal in evolutionary genetics. However, hybrid crosses between species in mammals often suffer from hybrid sterility, greatly complicating genetic dissection of trait variation. Here we describe a simple, robust and transgene-free technique to make “*in vitro* crosses” in hybrid mouse embryonic stem cells by inducing random mitotic crossovers with the drug ML216, which inhibits *Bloom syndrome* (BLM). Starting with an interspecific hybrid (between *Mus musculus* and *Mus spretus*) embryonic stem cell line spanning 1.5 million years of divergence, we demonstrate the feasibility of mapping enzymatic differences across species within weeks and the possibility of re-deriving whole mice. Our work shows how *in vitro* crosses can overcome major bottlenecks like hybrid sterility in traditional mouse breeding to address fundamental questions in evolutionary biology.

**Impact Statement:** By mixing hybrid mouse genomes in stem cells via mitotic recombination, genetic mapping and hybrid mosaic mice can be achieved in weeks, even across species barriers.

---

Discovering the genetic changes underlying species differences is a central goal in evolutionary genetic (1). However, hybrid crosses between even recently diverged species in animals often suffer from hybrid sterility (1, 2), greatly complicating genetic mapping of trait variation, especially in mammals. On the other hand, within-species genetic mapping has been tremendously successful in linking genetic polymorphisms to trait variations in innumerable organisms since the early twentieth century (3-5). Almost all mapping studies across diverse species have depended on meiotic linkage mapping panels generated through breeding to identify genetic loci controlling trait variations, or quantitative trait loci (QTL). Mapping resolution depends largely on crossovers arising from meiotic recombination to disentangle linked genetic associations. Accordingly, to achieve high-resolution mapping to the level of individual genes, researchers are driven to create ever-larger mapping populations and/or accumulating recombination over at least two, often many generations (6-8). In this respect, genetic studies in the mouse are complicated by the relatively long generation times and small litter sizes, which often decline further over generations due to increased inbreeding. Consequently, compared to yeast, worms and *Arabidopsis* (6-8), genetic mapping in the mouse requires far greater resources, yet relatively few traits have been mapped to the gene level (but see landmark studies identifying *Tlr4* and *Prdm9*) (9, 10). This challenge was particularly acute for panels involving divergence at or beyond the species level, where the difficulty or impossibility in generating fertile crosses calls into question whether the panel could be generated in the first place. Nonetheless, the potential to reveal unique biology occurring at the species boundaries in mammalian evolution makes such panels worthy attempts, even allowing for lower mapping resolution (11-15). This is because evolutionary changes in trait architecture can reveal much about the underlying evolutionary process. In this respect, direct assaying of hybrid genomes offers advantages unmatched by simpler single-gene functional assays or comparative transcriptome or sequence analyses, because it integrates actual interactions of every gene in the hybrid genomes. We argue that even cellular or expression phenotypes from such recombinant hybrids should offer unique insight into genome function and evolution. Should genetic exchange in hybrid animal genomes become feasible, direct genetic mapping of species differences would become routinely possible.

We set out to establish a universal method that allows genetic mapping in mammals without breeding, even across divergent species. We choose to initially focus on a cellular system based on mouse ES cells, which opens up the possibility of employing the full range of genetic manipulation available in tissue culture systems. We also anticipate that the rise of national biobank repositories for human induced pluripotent stem cells (iPSCs), together with the rapid development of organoid assays will ultimately counteract current limits in a purely cellular phenotyping system. In fact, “cellular phenotyping” offers many advantages over organismal assays in scale, costs and reproducibility. Therefore we have chosen hybrid mouse ES cells as an ideal setting to establish an *in vitro* cross system. A minimal system will have the two following features: an ability to induce on-demand extensive genetic exchange; and genetic (and trait) variation such as those found in F1 hybrid ES cells.

Intriguingly, the technique to create genetic variation through recombination has been in broad use in the mouse genetics community, albeit never explicitly in F1 hybrid ES cells with the goal of genetic mapping. In 2004, two independent groups showed that recessive, biallelic mutants could be reliably recovered in mouse ES cells without breeding by suppressing *Bloom Syndrome* (BLM; Fig. 1a) (16, 17). Yusa and coworkers showed that these recessive mutants arose via mitotic recombination between homologous chromosomes (18). We reasoned that the same mechanism could be leveraged to generate genome-wide random *mitotic recombination*. This mechanism enabled the creation of panels of arbitrary size carrying recombinant genomes, while avoiding the limitations of hybrid sterility or inbreeding depression (Fig. 1b).

**Fig. 1.**
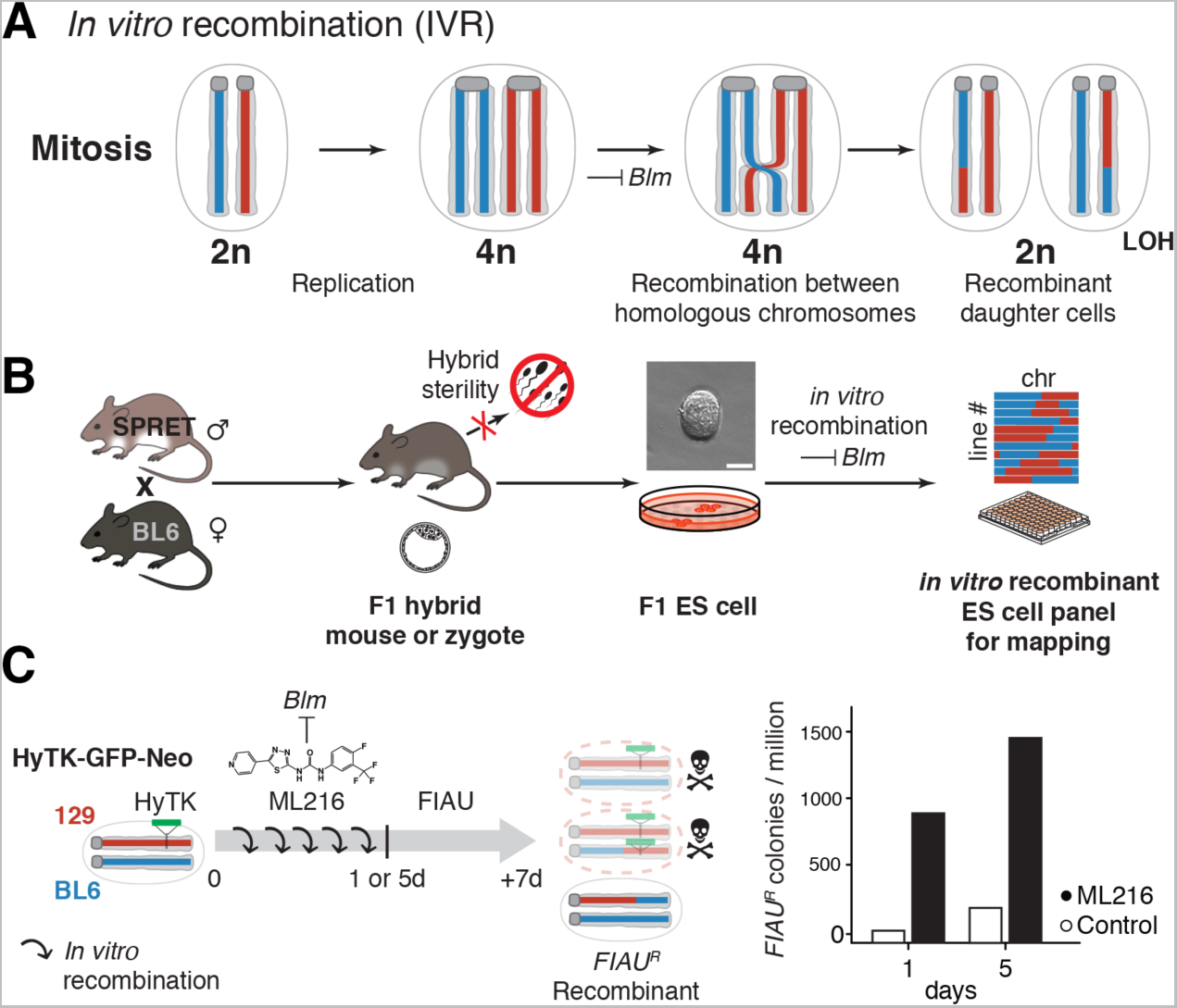
*In vitro* recombination via *Bloom syndrome* suppression. **(A)** *Bloom syndrome* (*Blm*) encodes a helicase normally active during mitosis. Loss of *Blm* activity leads to increased improper sister chromatid exchange as well as recombination between homologous chromosomes. Mitotic recombination can give rise to recombinant diploid daughter cells with loss of heterozygosity (LOH) between the breakpoint and the telomeres. **(B)** *In vitro* recombination (IVR) allowed the circumvention of hybrid sterility in crosses between the laboratory mouse, e.g., C57BL/6J (BL6) and a murine sister species *Mus spretus* (*SPRET*). (BL6 x *SPRET*)F1 hybrid mice were viable and allowed derivation of F1 ES cells despite male sterility (19). Applying IVR to F1 ES cells allowed rapid and efficient generation of recombinant ES cell panels for genetic mapping. Scale bar = 50 μm. **(C)** Efficiency of IVR was estimated by colony survival assay. We estimated recombination rate between homologous chromosomes with cells hemizygous for a dominant selectable marker (hygromycin phosphotransferase-thymidine kinase, abbreviated HyTK, green). We induced IVR by adding a small molecule BLM inhibitor ML216 (20) to the culturing medium for 1 or 5 days (d). Under fialuridine (FIAU) negative selection, cells having undergone mitotic recombination to become homozygous for the wildtype BL6 alleles (blue) survived; while non-recombined cells or recombinant cells retaining the HyTK transgene metabolized FIAU, resulting in cell death due to misincorporation of toxic nucleotide analogues (top and middle cells with red chromosomes). Under ML216 treatment (25 *μ*M), IVR rate was estimated to be 2.9×10^−4^ per cell per generation, yielding 800– 1500 FIAU-resistant colonies per million following treatment.

To test if BLM inhibition could lead to elevated homologous recombination rates in mitosis, we inhibited BLM in a number of mouse ES cell lines using a recently discovered small molecule inhibitor ML216 (Fig. 1c) (20). As a first test, we started with F1 ES cells between two laboratory mouse strains (C57BL/6J and 129, abbreviated to “BL6” and “129” here) that carried a targeted transgene as a hemizygous allele at the *GtRosa26* locus on distal Chromosome 6. We estimated homologous recombination by counting colony survival under fialuridine (FIAU) treatment, which selected against the transgene, which carried the dominant marker hygromycin phosphotransferase–thymidine kinase (HyTK) and green fluorescent protein (GFP; Fig. 1–Figure Supplement 1). We found that BLM inhibition led to highly elevated rates of homologous recombination, as revealed by increased numbers of FIAU-resistant colonies (Fig. 1c; *in vitro* recombination rate: 2.9×10^−4^ per cell per generation) and the appearance of mosaic GFP expression within a colony (Fig. 2a, right panels). This is broadly consistent with previously reported rates under direct *Blm* suppression or disruption (targeted tetracycline inhibition or knockout alleles: 2.3–4.2×10^−4^; compared to wildtype rates between 8.5×10^−6^–2.3×10^−5^) (16, 17). The small molecule BLM inhibitor ML216 offers unique experimental advantages, because its application is simple, rapid and reversible, eliminating the use of transgenes for *Blm* disruption or suppression (16, 17) or repeated transfections of small interfering RNA to achieve continued suppression of *Blm*. Importantly, elevated homologous recombination under BLM inhibition is not associated with increased aneuploidy (n=154 metaphase spreads; Mann-Whitney U test, W=1871, *h_1_*>0, n.s.; Fig. 1–Figure Supplement 2a). Further, ML216-treated ES cells retained robust expression of NANOG, a key marker for stemness (Fig. 1–Figure Supplement 3).

**Fig. 2.**
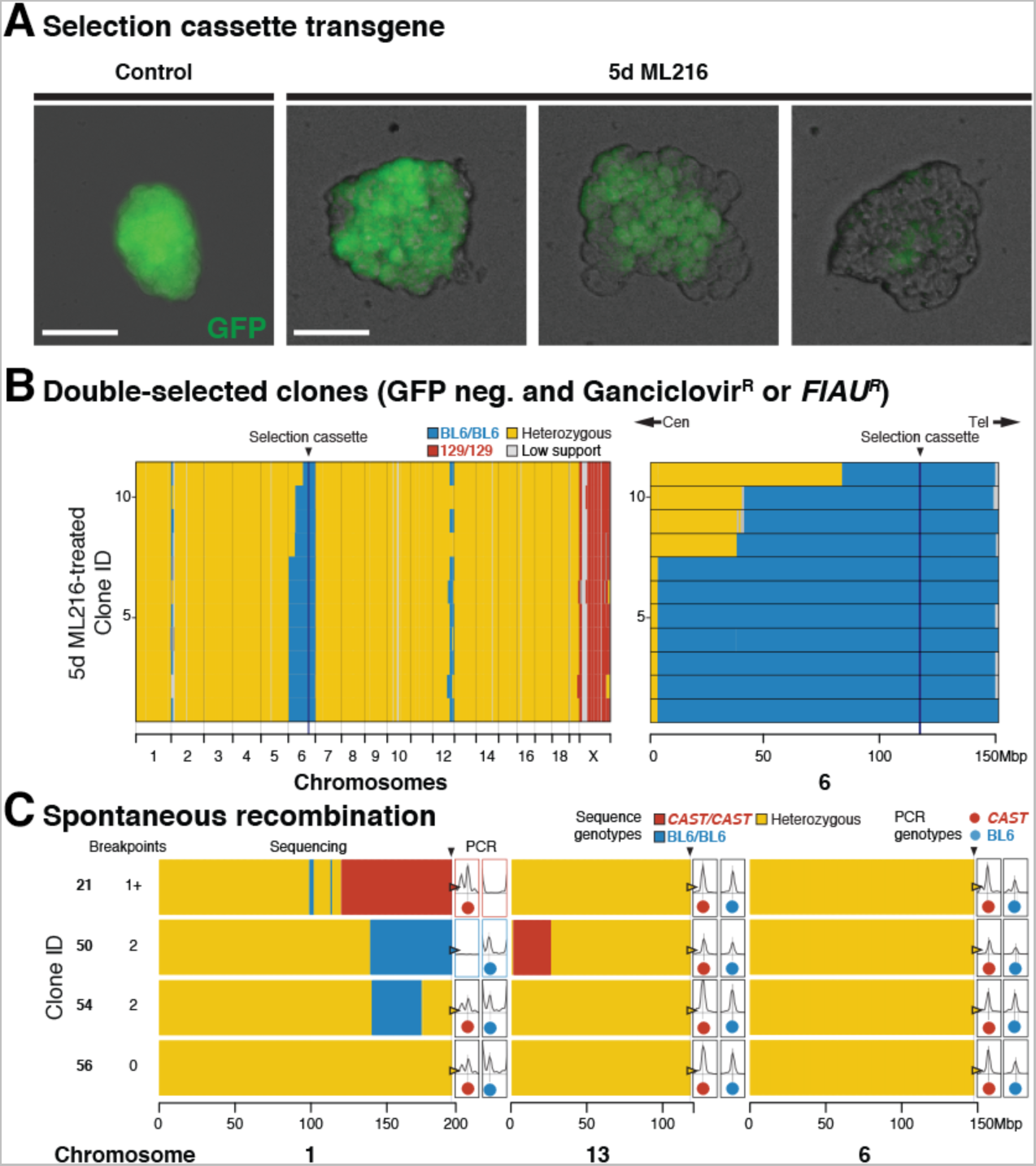
Widespread *in vitro* recombination across a range of evolutionary divergence. **(A)** ES cell colonies displayed mosaic GFP expression within a colony when cultured with ML216, but not under control conditions, consistent with homologous recombination and loss of GFP through IVR. Recombination between homologous chromosomes could result in daughter cells with two wildtype (BL6 allele, green) or transgenic copies (129 allele, not green). Early recombination events followed by random cell loss during clonal expansion could produce completely dark colonies. Scale bar = 100 μm. **(B)** After expansion under negative selection against the transgene (both ganciclovir and FIAU kill cells expressing HyTK), 11 ganciclovir-resistant and GFP-negative colonies were whole genome sequenced. Selection favoured loss of transgene (homozygous BL6/BL6 genotypes) at distal Chromosome 6. In contrast to normal meiotic recombination (averaging 1 or more crossovers per chromosome pair), mitotic recombination typically affected only a single chromosome pair: much of the genome remained heterozygous (yellow), with the exception of the transgene-carrying chromosome 6 (mostly BL6/BL6, blue) and the single 129 Chromosome X (male, 129, red). Mitotic recombination events converted genotypes telomeric to the breakpoint towards homozygosity (LOH, yellow to blue). **(C)** IVR also occurred in cells carrying divergent genomes with no transgenes. (BL6 x *CAST*)F1 hybrid ES cells were treated with ML216 and screened by PCR genotyping at diagnostic telomeric markers. Selected clones (two recombinant and control clones each) were whole genome sequenced, showing recombination events towards both homozygous genotypes, consistent with PCR genotype screening results (total breakpoints per clone ranged from 0–2). Additional recombination events were also recovered, even though the Chromosome 1 telomeric marker remained heterozygous (clone 54). These clones also carried non-recombined chromosomes (e.g., Chromosome 6, fully heterozygous, yellow).

To determine the frequency and distribution of mitotic crossovers under ML216-mediated BLM inhibition, we sequenced and compared the genomes of 11 clones that survived ganciclovir selection (a FIAU alternative; Fig. 2b). We also treated F1 hybrid ES cells derived from BL6 and *Mus castaneus* (diverged ~1 million years ago; CAST/EiJ, abbreviated to *CAST* here) (21) with ML216 but otherwise grown under non-selective conditions. Using the transgene-free (BL6 x *CAST*)F1 line (21), we screened 46 randomly-picked ML216-treated clones for spontaneous LOH recombinants and recovered recombinants in both directions on Chromosome 1. Sequencing of specific recombinant clones revealed conversion from F1 heterozygous genotypes towards both BL6/BL6 and *CAST*/*CAST* homozygous genotypes at the telomeres (Fig. 2c, clones 21 and 50, note also additional recombination on Chromosome 13). In contrast, control non-recombinant clones retained heterozygosity at the telomeres (clone 54 and 56). But even here we discovered a single clone carrying additional internal recombinants on Chromosome 1 (Fig. 2c).

Genome-wide sequencing of the recombinants revealed several striking patterns. First, crossover breakpoints were distributed along the entire chromosome (e.g., Chromosome 6 in Fig. 2b), echoing previous reports in other *Blm*-suppressed ES cells (18), suggesting recombinants can be used for mapping this as well as other cellular traits. Second, FIAU selection strongly and significantly enriched for recombinant chromosomes compared to unselected conditions (n=11 out of 11 vs. 9 out of 826; Fisher Exact Test, *P <* 2.2×10^−16^), with the recombination map biased strongly by the location of the selection cassette (all 11 crossovers were centromeric to Chromosome 6, 113Mbp, Fig. 2b). Our data suggest that chromosome segments telomeric to the cassette did not affect selection and were free to recombine. This observation, while relatively trivial for a targeted transgene here, was nonetheless instructive in the following, more nuanced cases involving natural variations. Third, crossovers created by mitotic recombination usually occurred only on one or few chromosomes at a time (Fig. 2b, c; Fig. 2–Figure Supplement 1), in stark contrast to an average of one crossover per chromosome arm during meiosis. Importantly, our results are in broad agreement with other reports of mitotic recombination (18, 22). Taken together, the data show that BLM inhibition efficiently generated *in vitro* recombination (IVR) across wide evolutionary distance and IVR ES cell panels may constitute genetically distinct lineages ideal for genetic mapping.

Our experiments to determine IVR rate demonstrated that among the chromosome-wide recombination positions their collective location and LOH state were indicative of the position of the selectable transgene (HyTK or GFP), with the major difference being that under mitotic recombination or IVR, the critical interval was defined only on the centromeric side. While it would have been trivial to screen for additional lines to increase mapping resolution towards *GtRosa26*, we chose to further illustrate the potential of this approach by mapping naturally-occurring variations using IVR. One classical polymorphism is the 25 to 75-fold increased activity of the *Mus spretus* “a” allele of hypoxanthine-guanine phosphoribosyltransferase (*Hprt^a^*) compared to the laboratory mouse *Hprt^b^* allele (23). Importantly, HPRT metabolizes the anti-metabolite tioguanine (6-TG) and causes cytotoxicity. It should be noted that beside the known *Hprt* polymorphism, tioguanine susceptibility itself has not been previously mapped genetically within or between mouse species. Here, we expected ES cells carrying *Hprt^a^* to be highly susceptible to 6-TG treatment, whereas *Hprt^b/b^* or *Hprt^-/-^* ES cells should survive far higher 6-TG concentrations (Fig. 3–Figure Supplement 1). We set out to map the QTL for differential 6-TG susceptibility using a bulk segregant assay simply by comparing allele frequencies across the genome between pools of 6-TG susceptible and resistant ES cells as determined by flow cytometry. We called this procedure “flow mapping”, which takes advantage of a structured but genetically diverse population of cells in tissue culture (but also see “X-QTL” in yeast, {Ehrenreich 2010}).

We first confirmed the absence of chromosome-scale rearrangements between the parental strains that could preclude mapping using the *de novo* assembled genomes of the parental strains made available by the Wellcome Trust Sanger Institute (BL6 and SPRET/EiJ, abbreviated to *SPRET* here) (25, 26). We generated IVR panels by treating a female (BL6 x *SPRET*)F1 hybrid ES cell line (“S18”) (19) with ML216 over 5, 10 and 21 days (d; Fig. 3a). The use of a female ES cell line, which carried two active X chromosomes prior to the onset of X inactivation during differentiation (27), allowed direct selection on the alternative *Hprt*^a^ and *Hprt^b^* alleles. After confirming biallelic *Hprt* expression in S18 cells using quantitative PCR, we treated control and IVR S18 cells with 6-TG and determined cell viability via a 4′,6-diamidino-2-phenylindole (DAPI) exclusion assay. Damaged cells with ruptured membrane exhibited rapid uptake of DAPI, a feature unaffected by ML216 treatment, and were distinguishable by fluorescent-activated cell sorting (FACS; “Live” proportions under ML216 treatment vs. “Live” proportions under 6-TG treatment, n = 5 paired treatments; Kruskal-Wallis test, *χ*^2^ = 13.17,d.f.= 1, *P <* 0.0003; Fig. 3a; Fig. 3–Figure Supplement 2). We separately recovered and sequenced each “Resistant” (*6-TG^R^*) and “Susceptible” (*6-TG^S^*) pool (Fig. 3a). Under both 5d and 21 d ML216 treatment, a large skew towards enriched *SPRET* coverage was observed on Chromosome X in the *6-TG^S^* relative to the *6-TG^R^* pool (Fig. 3a, b). This was in stark contrast to the genomic background, which showed little deviation from equal *SPRET* and BL6 contributions (normalized *SPRET* coverage for Chromosome X: 1.10, 95% confidence interval: 1.02–1.19; autosomes: 0.998, conf. int.: 0.986–1.01). The genome-wide peak *SPRET* enrichment window was found on Chromosome X, and it contained the *Hprt* gene itself (normalized *SPRET* coverage in *6-TG^S^* pool, 1 Mbp window: 1.19, conf. int.: 1.09–1.28). Our results are consistent with the known role of *Hprt* in mediating 6-TG susceptibility and thus the gene underlying the QTL on Chromosome X. Here, our results through forward genetic mapping for 6-TG susceptibility clearly identified a single locus, providing strong evidence that 6-TG susceptibility depended only on *Hprt* genotypes. While flow mapping combined superior mapping resolution and experimental simplicity as a bulk experiment, recovery of individual clones could provide further proof through the recovery of specific recombination breakpoints confirming the role of *Hprt* in mediating differential 6-TG susceptibility and exclusion of alternative modes of resistance such as aneuploidy. To recover specific recombinant breakpoints, we also sequenced 46 individual *6-TG^R^* IVR clones after 10d ML216 treatment (Fig. 3c). Echoing the skewed crossovers patterns centromeric to the HyTK selection cassette (Fig. 2b), we observed more *SPRET*-to-BL6 than BL6-to-*SPRET* recombinants (35 vs. 8, *P* ≤ 2 × 10^−5^, exact binomial test, *h_1_*≥ *h_0_*). We note, however, that despite the strongly skewed ratio of 27 BL6/BL6 homozygous clones at the *Hprt* locus out of 46 total recovered clones, we still observed 9 heterozygotes and 10 *SPRET*/*SPRET* homozygous clones (BL6/BL6 58.6%; Chi-squared test using observed allele frequencies, *χ*^2^ = 13.17, d.f. = 2, *P* ≤ 0.002). This could be due to a quantitative, rather than absolute allelic difference in susceptibility to 25μM 6-TG treatment (Fig. 3–Figure Supplement 1); non-exclusive FACS gating based on DAPI exclusion; or other new mutation(s) at *Hprt* or elsewhere leading to 6-TG resistance (16). Taken together, we conclude that we were able to perform forward genetic mapping using IVR and recover *Hprt* as the gene underlying 6-TG susceptibility differences between BL6 and *SPRET*.

**Fig. 3.**
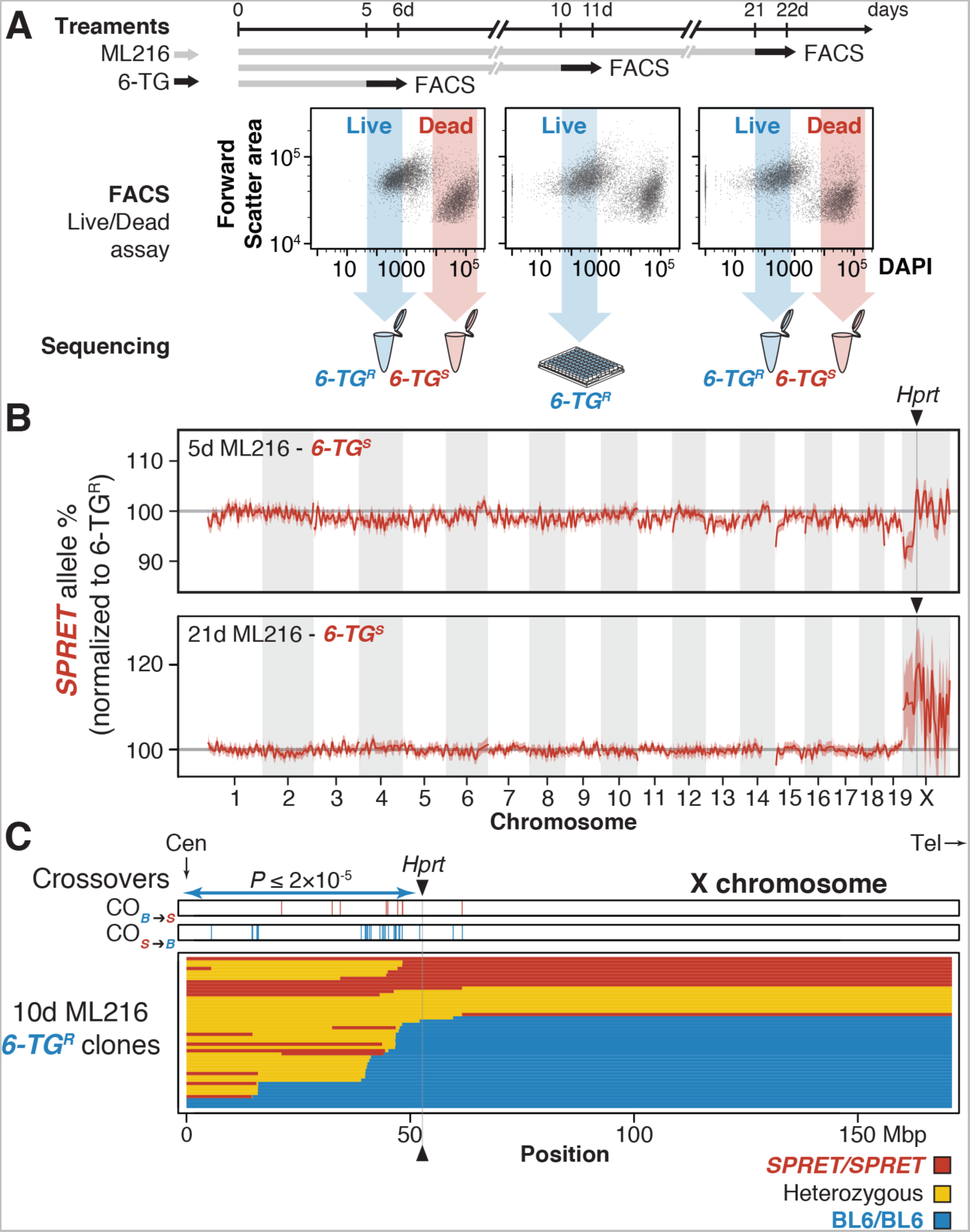
*In vitro* genetic mapping of variation in tioguanine susceptibility between divergent species. **(A)** A female ES cell line S18 derived from a *Mus spretus* and C57BL/6N F1 interspecific hybrid was treated with ML216 (25 μM) and subjected to the anti-metabolite tioguanine (6-TG) for 1d prior to fluorescence-activated cell sorting (FACS). ES cells were evaluated for viability based on 4′,6-diamidino-2-phenylindole (DAPI) exclusion. Resistant and susceptible (*6-TG^R^* and *6-TG^S^*) sub-populations were gated conservatively (shaded arrows) and pooled for sequencing. Individual clones from the 10d ML216 treatment were cultured and whole genome sequenced. **(B)** Skewed allelic contributions between the *6-TG^R^* and *6-TG^S^* pools suggested that the *SPRET* allele on Chromosome X conferred 6-TG susceptibility. Allele frequencies were normalized against *6-TG^R^* sample as an internal ML216 treatment control. Plotted are per megabase mean *SPRET* allele frequencies ± s.e.m. after 5 d and 21 d ML216 treatment. In both cases, the genome-wide peak window contains the *Hprt* gene with the *SPRET* allele showing significantly increased susceptibility. **(C)** Individual *6-TG^R^* clones following 10 d ML216 treatment were sequenced to determine recombination breakpoints. Crossovers in clones surviving 6-TG treatment recombined significantly more likely in the *SPRET*-to-BL6 direction (*S>B* = 37; *B>S* = 5; *P≤* 2×10^−5^) between the centromere and *Hprt*, consistent with strong selection favouring the BL6 *Hprt^b^* allele. In contrast, only 3 additional crossovers were detected telomeric to *Hprt*. At *Hprt*, most 6-TG surviving clones are homozygous for the *Hprt^b^* allele (27 vs. 9 heterozygotes and 10 *Hprt^a^* homozygotes).

The ability to easily circumvent hybrid sterility in evolutionarily divergent murine species led us to ask what developmental phenotypes may arise from such otherwise inaccessible genetic configurations (*M. spretus–*laboratory mouse hybrid males are sterile, following Haldane’s rule). Backcrosses using female hybrids are possible but extremely challenging)(15). Assaying developmental phenotypes from evolutionarily divergent hybrid ES cells is non-trivial, because hybrid sterility blocks germ line transmission. Since conventional re-derivation of whole mice from ES cells depends on germ line transmission through an intermediate chimera generation, alternative methods that directly generate fully ES cell-derived mice would have to be used. Accordingly, we produced fully ES cell-derived founder animals using laser-assisted morula injection (28) with two karyotypically normal but genetically distinct IVR ES cell lines along with the reference, non-recombined S18 cells (IVR 1 and 2; Fig. 1–Figure Supplement 2; Fig. 4–Figure Supplement 1; Movies 1–3). We succeeded in obtaining multiple embryos per line at embryonic (E) 14.5d of development (n=36, 24 from IVR lines vs. n=9 untreated S18 line). Using high-resolution micro-computer tomography (microCT), we observed that the embryos from the untreated clones showed uniformly normal development, whereas embryos from both IVR lines ranged from showing normal development to dramatic craniofacial and neural tube closure defects (2 abnormal embryos out of 4 scanned embryos in IVR line 1; 2 out of 7 in line 2; and 0 out of 6 from the original S18 line; Fig. 4; Fig. 4–Figure Supplement 2; Movies 1–3).Neural tube and craniofacial defects are among the most common developmental defects due to the complex coordination of cell migration and cell–cell communications, which may be impaired due to novel genetic interactions between homozygosed loci in the IVR lines (Fig. 4–Figure Supplement 1). Future studies will focus on establishing firm genetic links to such defects. Besides major developmental defects, we also identified and obtain 3D measurements from specific organs, including sub-regions of the brain, the heart and the liver, in multiple individuals from each ES cell line. Given an expanded panel of IVR ES lines, this approach illustrates the potential of characterizing, or even mapping, the genetic basis of evolutionary developmental variation. Despite the small sample size, our results show for the first time the feasibility and exciting opportunity to quantify and genetically map variation in developmental phenotypes in mammals using recombinants from evolutionarily divergent species.

**Figure 4.**
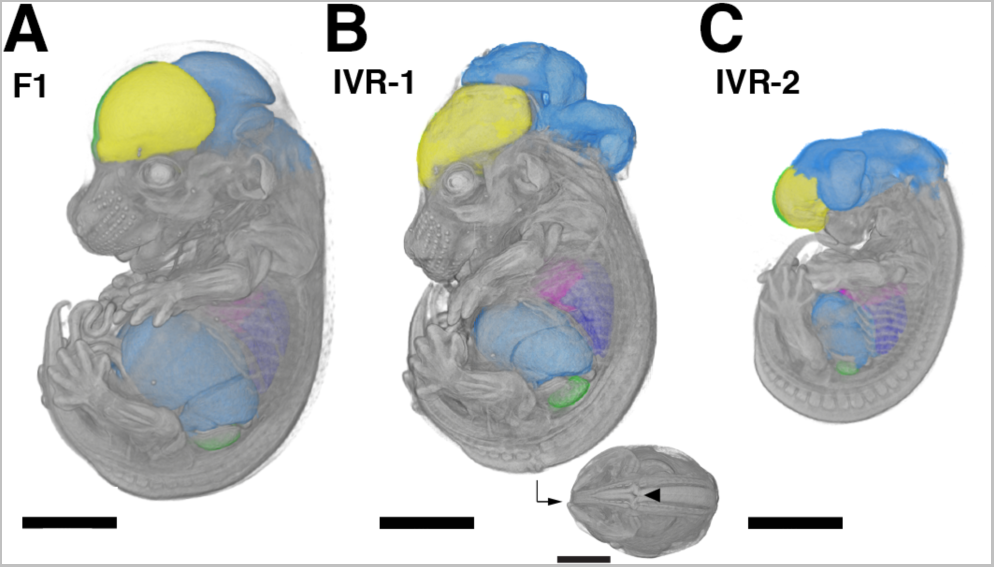
Accessing developmental phenotypes in recombinants between evolutionarily divergent species. Embryos at mid-gestation (14.5 d after fertilization) with nearly exclusive ES cell contribution were derived from non-recombinant F1 S18 ES cells **(A)** and IVR lines 1 **(B)** and 2 **(C).** Embryos were dissected, contrast-stained and scanned using X-ray micro-computer tomography at 9.4 μm resolution. The high scanning resolution allowed identification and precise measurements of individual organs (colorized here). Major developmental craniofacial and neural tube closure defects were observed in the IVR lines (**B**, caudal view with arrowhead indicates neural tube lesion). Scale bar = 200 μm.

A central goal of evolutionary genetics is to identify how mutations arose during evolution and influenced phenotypes. For many organisms, a major barrier has been the inability to reliably generate diverse and large mapping panels of sufficient evolutionary diversity. Here we describe a simple and robust method to make “*in vitro* crosses”, resulting in functionally intercross panels from otherwise sterile interspecific hybrid crosses. Being able to bring forth genetic diversity in a petri dish creates the unique opportunity to conduct mouse genetic mapping at unprecedented speeds with “flow mapping” (similar to “X-QTL” in yeast) (24) or arbitrarily large panels unmatched by most other model organisms, except possibly yeast (22, 24). As indefinitely renewable stem cell lines, IVR panels can be expanded, archived and shared, offering an arguably more feasible and cost-effective platform with many of the advantages sought from traditional community resources such as recombinant inbred line panels. Recently, Sadhu and coworkers have also achieved a major advance in genetic mapping using CRISPR/*Cas9*-mediated mitotic recombination in yeast (22). In contrast to CRISPR targeting, our transgene-free approach offers the simplicity of inducing genome-wide recombinants by the simple addition of a single inexpensive small molecule to the tissue culture medium. Further, we have shown that our IVR method works in a broad range of ES cells. With millions of potentially recombinant (thus genetically distinct) ES cells in a petri dish, we demonstrated how IVR enabled us to map a QTL for drug resistance in as few as 6 days (with an estimated total of 5 doublings over 5 days under ML216 treatment). Putting this in context, such an experiment using traditional mouse crosses would have taken 450 days, based on the typical mouse generation time of 90 days, assuming that hybrid sterility could be overcome and allowing for selfing. Going forward, we envision a combined, complementary approach to IVR: using BLM inhibition for mapping panel generations and efficient QTL identification, then switching to targeted transgene-based screening or CRISPR/*Cas9*-based IVR for fine-scale mapping.

As a novel mapping system, we observed a number of key differences between IVR and conventional meiotic genetic mapping. First, loss-of-heterozygosity due to mitotic recombination tend to occur between the breakpoint and the telomeres. Unlike conventional breeding with random assortment, under IVR in F1 hybrids, outcrosses are not possible. As a result, we tend to observe only heterozygous genotypes near the centromere, with informative crossovers almost always found between the centromere and a selectable QTL but not on the telomeric side. This asymmetry often led to a plateau in the association profiles from the QTL towards the telomeres on a given chromosome (Fig. 2b and 3c), an effect also seen in (22). As a consequence, interval mapping in IVR analogous to those in meiotic panels yields excellent genetic resolution on the centromeric side but poor resolution on the telomeric side [see distribution of crossover directions and breakpoints in Fig. 3c and (22)]. Second, access to common tissue culture methods under IVR greatly mitigates typical concerns such as panel sizes and power calculation in generating meiotic mapping panels. Since it is trivial to freeze samples and introduce selectable markers at any given locus or targeted chromosome breaks with a Crispr/Cas9 panel (22) with ES cells under tissue culture conditions, refinement of mapping resolution under IVR no longer depends on the diminishing return of breeding and screening for increasingly rare informative recombinants. To underscore this point, our flow mapping experiment for 6-TG susceptibility achieved mapping resolutions within 10 Mbp within weeks. Third, while it is true that mitotic recombination as used in IVR depends on error-prone repair of double-strand breaks that could affect phenotype through chromosome rearrangements and new mutations at breakpoints, two observations from our results may moderate this concern. One, we did not observe elevated aneuploidy under ML216 treatment, suggesting that IVR did not elevate rates of chromosome rearrangements (Fig. 1–Figure Supplement 2). Two, millions of variants already exist in our (BL6 x *CAST*) F1 or (BL6 x *SPRET*) F1 lines. These variants vastly outnumber any new mutations generated through IVR. Assuming a typical spectrum of mutation effects, these parental variants likely would contribute far more to trait variance than new mutations arising in a specific line. Since genetic mapping depends on testing for different genotypic effect of an allele across all lines carrying the same genotype at loci that are typically megabases away from a random double-strand breakpoint, it is reasonable to expect that the mutagenic effect of mitotic recombination should have a rather limited impact on genetic mapping. Under the flow mapping design, the mutagenic effect of mitotic recombination is further diluted, because millions, if not tens of millions of cells in bulk population cultures are phenotyped and sequenced as pools. This conclusion is supported by our ability to locate and map various transgenes or QTLs in this current study. We are nonetheless in the process to formally characterize the relative contribution to trait variation due to the mutagenic effects of IVR and that of the parental genomes.

Rather than thinking of IVR as a replacement of current genetic mapping methods, we see its establishment as an important extension to the existing toolkit that is complementary to whole organism methods. In the mouse, the largest organismal recombinant inbred (RI) panel BXD contains “only” ~160 lines (with most published work based on the ~35 original BXD strains)(29) and attempts in generating panels incorporating greater divergences encountered enormous challenges (30). Nevertheless, mouse RI resources still represented some of the most powerful tools available to dissect system genetics in the mouse, the prime biomedical model organism (31). Seen in this light, the mouse community should encourage alternative approaches that could greatly extend the available renewable resources, even in a cellular or tissue context, not least because the genotype combinations between divergent species would have been hitherto impossible to obtain in the first place.

Given its potential for broad applications, we are currently making improvements to many aspects of *in vitro* recombination to make trait mapping of interspecific differences in mouse or other mammals routine. This includes improving the efficiency of IVR panel creation from hybrid cell lines and developing robust phenotyping protocols beyond the proof-of-concept experiments we have presented here. For example, we have already made refinements in IVR to screen for lines carrying recombination on multiple chromosomes (to be published separately). We are also performing detailed characterization on the location and distribution of recombination breakpoints to determine if certain genome features promote mitotic recombination (see possibly clustered breakpoints in Fig. 2b & 3c). In addition to the traits we have investigated, *Mus spretus* and the *Mus musculus* laboratory mouse differ in a number of distinct traits. Comprehensive genetic dissection of traits such as longevity and telomere lengths (32), cancer and inflammation resistance (33, 34) and metabolism (35), would be extremely useful. Many of these traits have tissue or cellular models that can be used in the context of IVR. For cellular traits that can be measured as fluorescent signals, either from immunohistochemistry staining of marker proteins or antigens, flow mapping would be readily applicable. Especially in the area of stem cell biology and genome functional genomics, the IVR platform opens up entire new possibilities for exploring how evolutionary divergence affects genome functions through expression QTL mapping, ChIP or ATAC-Seq and where applicable, flow mapping (and further fine mapping using Crispr-directed recombination as appropriate). At this stage, testing of specific genes discovered through the process via knock-in gene replacement as whole mice would serve as validation of the *in vitro* findings. For derivation of whole mice from IVR lines, we acknowledge that our immediate attempt serves only to illustrate the possibility of obtaining whole animals from IVR genotypes, rather than a full-scale attempt at mapping developmental differences. It is important to stress that actual genetic mapping of developmental differences will require a much greater number of lines, ideally using improved, highly-recombinant IVR methods. Clearly, such a study will represent a major undertaking for single laboratories or small consortia. However, against the backdrop of systematic phenotyping performed on hundreds of lines in the mouse knock-out consortium, the costs and effort in screening a number of lines comparable to RI panels (around 35 lines) using IVR would not only be feasible, but also very likely to produce important results because it covers great evolutionary distances.

Future experiments may also probe even greater evolutionary divergence: early work has shown that F1 hybrids spanning as much as 6 million years between *Mus musculus* and *Mus caroli* was viable (36); or even human–mouse trans-chromosomal lines like *Tc1* (37). Further, given active development in single-cell genomics and disease modeling from patient-specific induced pluripotent stem cells (iPSC), especially with organoids or organ-on-a-chip microfluidics systems, we are optimistic that the *in vitro* recombinant platform can be broadly applied to mouse, human or even other species to accelerate the identification of the genetic basis of many traits and diseases.

## Acknowledgements

We thank Felicity Jones for experimental design, helpful discussion and input, and for improving the manuscript. We thank Caroline Schmid for animal husbandry. We thank Sebastian Kick for help with microCT scanning. We thank the Chan and Jones Labs members for support, insightful scientific discussion and improving the manuscript. We thank Derek Lundberg for help with library preparation automation. We thank Christa Lanz, Rebecca Schwab and Ilja Bezrukov for assistance with high-throughput sequencing and associated data processing; Andre Noll for high-performance computing support; Cornelia Grimmel and Stella Autenrieth for technical assistance with FACS; the MPI Tübingen I.T. team for computational support. We thank Rémi Blanc at FEI for assistance and support with 3D image analysis. RV-L3-HyTK-2L was a gift from Geoff Wahl (Addgene plasmid # 11684). pCAG-Flpo was a gift from Massimo Scanziani (Addgene plasmid # 60662). pBSII-IFP2-ORF was a gift from Nancy Craig. The AB2.2 ES cell line was a gift from Allan Bradley. The G4 ROSALUC ES cell line was a gift from Jody J. Haigh. We thank Hua Tang, David M. Kingsley, Karsten Borgwardt and Detlef Weigel for input and discussion on experimental design. J.P.L.C. is supported by the International Max Planck Research School “From Molecules to Organisms”. P.M. was supported by the Fulbright US Student Program. T.H. and C.L. are supported by Ghent University. Y.F.C. is supported by the Max Planck Society and a European Research Council Starting Grant #639096 “HybridMiX”.

## Author Contributions

Y.F.C. conceived the IVR strategy. M.N.C.F. and Y.F.C. designed the original experiments. S.L. and Y.F.C. developed the flow mapping strategy. S.L., and M.K. designed and perform the cell culture and sequencing experiments and analyzed the data. S.L. and J.P.L.C. performed cell culture experiments, screening for recombinants. P.M., M.N.C.F. and R.W. designed the original pilot experiments on the IVR strategy and performed the experiments. Y.F.C., S.L., and R.N. planned and performed the morula injection experiment and analyzed the data. J.G., T.H., C.L. provided critical ES cell lines. S.L., M.K., J.P.L.C., R.N., R.W., J.G., T.H., C.L. and Y.F.C. wrote the manuscript. All authors discussed the results and implications and commented on the manuscript at all stages.

## Materials and Methods

### Animal Care and Use

All experimental procedures described in this study have been approved by the applicable University institutional ethics committee for animal welfare at the Faculty of Sciences, Ghent University, Belgium, (reference number 06/022); or local competent authority: Landesdirektion Sachsen, Germany, permit number 24-9168.11-9/2012-5.

### Reference genome assembly

All co-ordinates in the mouse genome refer to *Mus musculus* reference mm10, which is derived from GRCm38. Sequence data have been deposited in the GEO database under accession number [X].

### Cell Culture

All ES cell lines used in this study are summarized in Table S1.

Unless otherwise stated, murine stem cell lines have been cultured on Attachment Factor Protein (AF) (ThermoFisherScientific, Schwerte, Germany) coated cell culture dishes on inactivated SNL 76/7-4 feeder cells (“feeder” plates; SCRC-1050, ATCC, Middlesex, United Kingdom) and using 2i/LIF media as follows: KnockOut Serum Replacement (ThermoFisherScientific), KnockOut DMEM (ThermoFisherScientific), 2-Mercaptoethanol, 1000x, 55 mM (ThermoFisherScientific); MEM Non-Essential Amino Acids Solution, 100x (ThermoFisherScientific); GlutaMAX Supplement, 100x (ThermoFisherScientific); 3 μM GSK-3 inhibitor CHIR99021 (Sigma–Aldrich, Munich, Germany); 1 μM MEK inhibitor PD0325901 (Sigma–Aldrich); insulin solution, human (Sigma–Aldrich), 1000 U/mL recombinant mouse LIF (expressed in-house).

Unless otherwise stated, cell culture media was replaced daily.

### BLM inhibition using ML216

BLM inhibition was performed using 25 μM ML216 (Sigma–Aldrich) in 2i/LIF media on inactivated feeders. Killing curve for ML216 was performed using the WST-1 assay (Roche, Basel, Switzerland) according to the manufacturer’s instructions.

### Plasmid construction

The pMK11 plasmid was constructed by blunt-end ligation of the pRMCE-DV1 plasmid’s backbone, after excision of its chloramphenicol–*ccdB* cassette between the EcoRV and SbfI sites, and replacing it with the HindIII-excised hygromycin phosphotransferase-thymidine kinase cassette (HyTK) from the RV-L3-HyTK-2L plasmid (38) (Plasmid # 11684, Addgene, Cambridge, USA). The final pMK11 construct contained flanking *FRT wt* and *FRT mutant* sites for recombinase-mediated cassette exchange detailed below.

### Generation of HyTK-EGFP-Neo cell line

G4 ROSALUC B12 ES cells (39) were co-transfected with pMK11 described above and *FLP* mRNA (StemMACS *Flp* Recombinase, Miltenyi Biotec, Bergisch Gladbach, Germany) or pCAG-Flpo (40) (Plasmid # 60662, Addgene) using Lipofectamine 2000 (ThermoFisherScientific). We replaced the cassette at the *GtRosa26* locus with a cassette carrying two selectable markers, HyTK and enhanced green fluorescent protein (EGFP, selectable in fluorescence-assisted cell sorting; Fig. 1–Figure Supplement 1). Successful replacement of the cassette with a re-activated neomycin resistance gene was selected for with 200 μg/mL geneticin (G148; ThermoFisherScientific). Resistant colonies were picked after 7 days (d) of selection and further expanded. Correct replacement was confirmed by junction PCR with primers SA_loxP_Rev: 5′–GCGGCCTCGACTCTACGATA–3’ and ROSA26_3HA_F_BamHI: 5′–GCGGGATCCCCTCGTGATCTGCAACTCC–3′. The presence of an intact BL6 wildtype allele was confirmed by an alternative reverse primer oIMR8545 5′–AAAGTCGCTCTGAGTTGTTA–3′. PCR was performed as a quantitative PCR reaction. See “RNA Extraction, Reverse Transcription and “Real Time PCR” section below for more details.

### Colony Survival Assay

HyTK-EGFP-Neo cells were seeded at a density of 5×10^5^ per 10 cm AF/feeder plate. Eight hours (h) following the plating, 25 μM ML216 treatment was initiated and continued for 1 or 5 d. Prior to the start of negative selection,cells were replated at 2×10 ^5^per 10cm AF/feeder plate and FIAU (0.2 μM, Sigma–Aldrich) or ganciclovir (10 μM, Sigma–Aldrich) selection was initiated after 1 d and continued for 5 d. In order to determine the plating efficiency after ML216 treatment, cells were plated at 1×10^3^ per 6 cm AF/feeder dish. Colonies were stained with the Alkaline phosphatase kit (EMD Millipore, Billerica, MA,USA). Before the application of negative selection, 20 random views of each plate were taken using an EVOS FL Cell Imaging System (ThermoFisherScientific) and counted using `Fiji v2. 0. 0-rc54/1.51h` (41).

### Screening for spontaneous recombinant ES cell colonies

Cells were plated at a density of 1×10^5^ per 3.5 cm AF plate. Treatment with 5 μM ML216 was initiated 16 h after plating, continued for 2 d and then followed by 3 d of 25 μM ML216 treatment. Cells were then re-plated on a 10 cm AF plate and cultured for 5 d in 2i/LIF without ML216. Two hundred colonies were randomly picked, and 153 were expanded for multiplexed genotyping.

### Multiplexed genotyping for detection of loss-of-heterozygosity (LOH)

Diagnostic insertions or deletions (indels) between BL6, *CAST* and *SPRET* strains that are greater than 20bp in length and located within the most distal 10Mbp of each chromosome were filtered from the publicly available variant panel from the Mouse Genomes Project made available by the Wellcome Trust Sanger Centre (`v5 dbSNP v142 release`) (19) using `VCFtools v0.1.14` (42). Automated primer design was carried out with Primer3 v.1.1.3 using the following parameters: PRIMER _OPT_SIZE=20; PRIMER _MIN_SIZE=18; PRIMER_PRODUCT_OPT_SIZE=300 PRIMER_PRODUCT_ SIZE _RANGE=250-400 PRIMER_MAX_SIZE=23 PRIMER_NUM_NS_ACCEPTED=1 PRIMER_LEFT_MIN_TM=58 PRIMER _LEFT_MAX_TM=62 PRIMER_RIGHT_MIN_TM=58 PRIMER_RIGHT_MAX_TM=62 PRIMER_MAX_DIFF TM=2 PRIMER_MIN_GC=45.0 PRIMER_MAX_GC=85.0 PRIMER_MAX_POLY_X=3 PRIMER_SELF _ ANY=4. Among indels with successfully designed primer pairs, the most telomeric amplicons were chosen, and an extension was added to either the forward (M13F) or reverse (M13R-46) oligonucleotide to allow for easy fluorophore incorporation as described in (43). The amplicon sizes were further optimized following pilot capillary sequencer runs to avoid amplicon size overlap in a multiplexed run. All primers and expected fragment sizes are listed in Supp. Table 2. For genotyping of cell colonies, primers pairs were pooled into 4 multiplexed PCR reactions. Group 1 (Chr1, Chr7, Chr13, Chr14 and Chr18) and Group 2 (Chr3, Chr6, Chr16, Chr17 and Chr19) primer mixes contained 2 and 4 μM of forward and reverse primers, respectively, for each listed chromosome plus 20 μM of the universal FAM-labeled M13F_FAM primer. Chromosomes 13 and 17 primers were mixed at 6 and 12 μM concentration. For Group 3 (Chr2, Chr4, Chr5, Chr11,ChrX) and Group 4 (Chr8, Chr9, Chr10, Chr12, Chr15) mixes, the forward and reverse primers were mixed at 4 and 2 μM concentration, along with the HEX-labelled M13R-46_HEX primer at a concentration of 20 μM. QIAGEN Multiplex PCR *Plus* Kit (Qiagen, Hilden, Germany) was used according to manufacturer’s recommendations (including the addition of 5× Q-Solution) at 10 μL final reaction volume with 3 to 10 ng of DNA per PCR reaction. The PCR program used was: 95 °C for 15 min, then 52 cycles of 94 °C for 30 s; Group-specific annealing temperature for 2.5 min; and 72 °C for 1 min; followed by a final extension at 72 °C for 30 min and hold at 4 °C. The group-specific annealing temperatures were: Group 1: 63 °C; Group 2: 63.8 °C; Group 3: 57 °C; and Group 4: 64 °C. Then the PCR reactions were pooled at equal 1 μL proportions and analyzed with a 3730xl DNA Analyzer capillary sequencer (ThermoFisher Scientific, Germany) using the fragment analysis program with the G5-RCT Dye Set. Electropherogram traces were analyzed with the Microsatellite Plugin in `Geneious v7.1.9` (44).

### 6-TG treatment and DAPI exclusion assay

Prior to the main experiments, killing curves for 6-tioguanine (Sigma–Aldrich) was performed using WST1 assay (Roche) according to the manufacturer’s instructions (Fig. 3–Figure Supplement 1). For the main experiments, S18 ES cell line was cultured for 5, 10 or 21 d with 25 μM ML216 prior the treatment with 25 μM 6-TG in 2i/LIF starting from an initial seeding concentration of 1 ×10^5^ cell per 3.5 cm AF plate. To avoid overcrowding, at day 3 of the ML216 treatment colonies were dissociated using Accutase Cell Dissociation Reagent (ThermoFisherScientific) and re-seeded on a 10 cm AF-plate while continuing ML216 treatment. At day 5, the cells were moved to a 15 cm AF plate prior to 6-TG treatment. After 16 h, 6-TG in 2i/LIF was added at a concentration of 25 μM. From each plate 2.5×10^4^ cells were used to continue the experiment until day 10 or 21. 4′,6-diamidino-2-phenylindole (DAPI) staining (1 μg/mL, Sigma–Aldrich) was employed for “live/dead” cell viability determination after 1 d of 25 μM 6-TG treatment. Briefly, ES cells were treated with ML216 and/or 6-TG to induce IVR and cell death, respectively. Colonies were dissociated using Accutase and re-suspended in phosphate buffered saline (PBS) within 1 h of analytical or preparative fluorescence-activated cell sorting (FACS). For details on FACS see below.

### Fluorescence-Activated Cell Sorting (FACS)

Flow cytometry was performed at the University Clinic <u>T</u>ü<u>bingen</u> Dermatology Clinic FACS Core Facility using an Aria II Cell Sorter (Becton Dickinson GmbH, Heidelberg, Germany). To determine cell viability, we performed the DAPI exclusion assay. After excluding cell aggregates, we defined the *6-TG^R^* and *6-TG^S^* populations using conservative interval gates based on evaluating the data from reference flow experiments with 6-TG-treated DAPI-stained ES cells. For cell population evaluations, flow cytometry data was exported from BD FACSDiva Software v8.0.1 (Becton Dickinson). We carried out basic data handling and log_10_ transformation using the R Bioconductor package flowCore (45). Since live and dead cells cluster also in other measurements, we took both forward scatter area (FSC-A) and DAPI into account for our quantification, rather than using a simple interval gate on the DAPI/Pacific Blue-A channel. We defined data-driven “Live” and “Dead” clusters using mclust v5.2 (46, 47) in 6-TG-treated experiments, considering ML216-treated and controls separately. We then classified each cell in to the “Live” and “Dead” clusters, applying a 5% uncertainty cut-off. “Live” and “Dead” proportions were then calculated from the confidently assigned cells. Data was visualized using the package flowViz (48) (Fig. 3–Figure Supplement 2).

### RNA Extraction, Reverse Transcription, and Real Time PCR

RNA was isolated using TRIzol Reagent (ThermoFisherScientific) with a single-step method following (49). Complementary DNA (cDNA) was generated using High-Capacity cDNA Reverse transcription kit (ThermoFisherScientific) with 500 ng of RNA per reaction according to the manufacturer’s instructions. The newly synthesized cDNA (20 μl reaction) was diluted 5 fold and quantitative PCR (qPCR) was performed with SYBR-select Master Mix for CFX (ThermoFisherScientific) using a CFX384 Real-Time PCR system instrument (BioRad, Munich, Germany). We used the following primers for allele-specific amplification and detection: *Hprt^a^* (*SPRET*) forward: 5′– CAAAGCCTAAGAGCATGAGCGC–3′, reverse: 5′-CAGAGGGAACTGATAGGCTGGC–3′, amplicon size: 229bp; *Hprt^b^* (BL6) forward: 5′–GCCAAATACAAAGCCTAAGATGAGCG–3′, reverse: 5′– CCAGCCTACCCTCTGGTAGATTG–3′, amplicon size: 236bp. The standard CFX mode for Tm ≥ 60 °C was used, with the following thermocycling program: 50 °C for 2 min, 95 °C for 2 min, followed by 35 cycles of 95 °C for 15 s, 60 °C for 1 min. Melting curve analysis over 80 steps of 0.5 °C increments was performed and curves inspected to ensure uniform annealing.

### Immunofluorescence staining

ES cells were cultured for 3 d on 12 mm cover glasses pre-coated with AF and feeder layer. Cells where then fixed 10 min in 4% paraformaldehyde, permeabilized 10 min in 0.25% Triton X, blocked in 5% serum for 1 h at room temperature. ES cell colonies were stained with anti-*Nanog* (1:100, rabbit, Cat # ab80892, Abcam, Cambridge, UK) antibodies for 2 h at room temperature and conjugated secondary antibody (1:400, anti-rabbit Alexa 467) for 1 h at room temperature. Nuclei were counter-stained for 5 min with DAPI at 1 μg/mL, mounted with ProLong Diamond Antifade Mountant (ThermoFisherScientific) and imaged using an AXIOVERT 200M inverted microscope (Zeiss, Oberkochen, Germany)

### Karyotyping

Metaphase spreads were prepared from Control and ML216-treated ES cells under 2i conditions (5 d culture for treatment on the original S18 background, 2 d for the IVR lines 1 and 2; see Cell Culture above for a detailed description of culturing conditions). Metaphase spreads were prepared essentially as described in (50) with the following modifications. Cells were initially plated at a density of 2×10^5^ cells per 10 cm AF-coated culture dish. Spreads were mounted with ProLong Diamond Antifade Mountant (ThermoFisherScientific). Metaphase chromosomes were imaged with a 63x objective in a Zeiss APOTOME AXIO Imager.Z1 (Zeiss) equipped with an Orca-flash4.0 digital camera (C11440-22CU, Hamamatsu, Herrsching am Ammersee, Germany) and coupled to HCImage v4.3.5.8 image acquisition software. Chromosomes were anonymized and independently counted twice manually in `Fiji v2.0.0-rc-54/1.51h using` the multi-point tool.

### Sequencing and analysis pipeline

Sequencing libraries for high-throughput sequencing were generated using Nextera DNA Library Prep Kit (Illumina, Inc., San Diego, USA) according to manufacturer’s recommendations or using equivalent *Tn5* transposase expressed in-house as previously described (51). Briefly, genomic DNA was extracted from FACS-sorted clones, single colonies or pooled samples by standard Protease K digestion (New England Biolabs GmbH, Frankfurt am Main, Germany) followed by AmpureXP bead (Beckman Coulter GmbH, Krefeld, Germany) purification. Extracted high-molecular weight DNA was “tagmented” by commercial or purified *Tn5*-transposase. Each tagmented DNA sample was then PCR amplified with Q5 High-Fidelity DNA Polymerase (New England Biolabs) using barcoded i7-index primer (N701-N763) and the N501 i5-index primer. Pooled libraries were sequenced by a HiSeq 3000 (Illumina) at the Genome Core Facility at the MPI Tübingen Campus. Sequenced data were processed using a custom pipeline consisting of data clean-up, mapping, base-calling and analysis based upon fastQC v0. 10.1 (52); trimmomatic v0.33 (53); bwa v0.7.10-r789 (54); GATK v3.4-0-gf196186 modules MarkDuplicates and IndelRealignment (55, 56); samtools v1.2 (57, 58); bcftools v1.2 (59); and R v 3.2.0 (60). Genotype calls were performed against known informative single and multiple nucleotide variants between C57BL/6NJ, CAST/EiJ and SPRET/EiJ strains made available by the Wellcome Trust Sanger Centre (Mouse Genomes Project version 3 dbSNP v137 release (25). Coverage depths for the reference and alternative alleles were calculated based on the DP4 field in the variant VCF files. For individual clones, crossover breakpoints were called by TIGER (61), using default parameters. Custom Perl scripts were used to process files prior to plotting and visualization in R. Scripts are available upon request.

### Laser-assisted morula injection

Fully ES cell-derived embryos were obtained essentially as previously described in (27). Briefly, female C57BL/6NCrl mice were mated and host embryos harvested. ES cells from untreated S18 line and two IVR lines were injected into 8-cell stage embryos (morulae) after perforation of the zona pellucida with a laser pulse. After incubation for 1.5–2 h, injected embryos were transferred into the oviducts of E0.5 pseudo-pregnant CD1-ICR female foster mice. The host mice were monitored for recovery and development. At 14 d after the embryo transfer (approximating developmental stage E14.5), the gestation was terminated, and embryos were individually dissected, fixed with 4% paraformaldehyde for 45 min and stored in PBS. All manipulations were performed by R.N. or under R.N.’s supervision at the Transgenic Core Facility at the Max Planck Institute of Cell Biology and Genetics, Dresden, Germany.

### Micro computer-tomography (microCT)

Prior to scanning, embryos were perfused for 4 d in 25% Lugol’s, or iodine-potassium iodide solution. Contrast-stained embryos were rehydrated and mounted in 1 % agarose and scanned with a Skyscan 1173 instrument (`Control software version 1.6, Build 15`; Bruker Corporation, Billerica, MA, USA) at 9.96 micron (μm) resolution using a 0.5 mm aluminium filter with energy settings at 70 kV and 110 μA. Volume reconstructions were performed using `NRecon v.1. 6. 10. 4` (Bruker Corporation) using parameters determined based on fine-tuning for each scan (misalignment correction: 23-30; beam-hardening correction: 25%; ring-artifact correction: 10; no smoothing). Image analysis, segmentation and visualizations were performed using `Amira v6.2.0` (FEI, Hillsboro, OR, USA) with the `XImagePAQ extension 6.2`.

## Figure Supplements

**Fig. 1–Figure Supplement 1.**
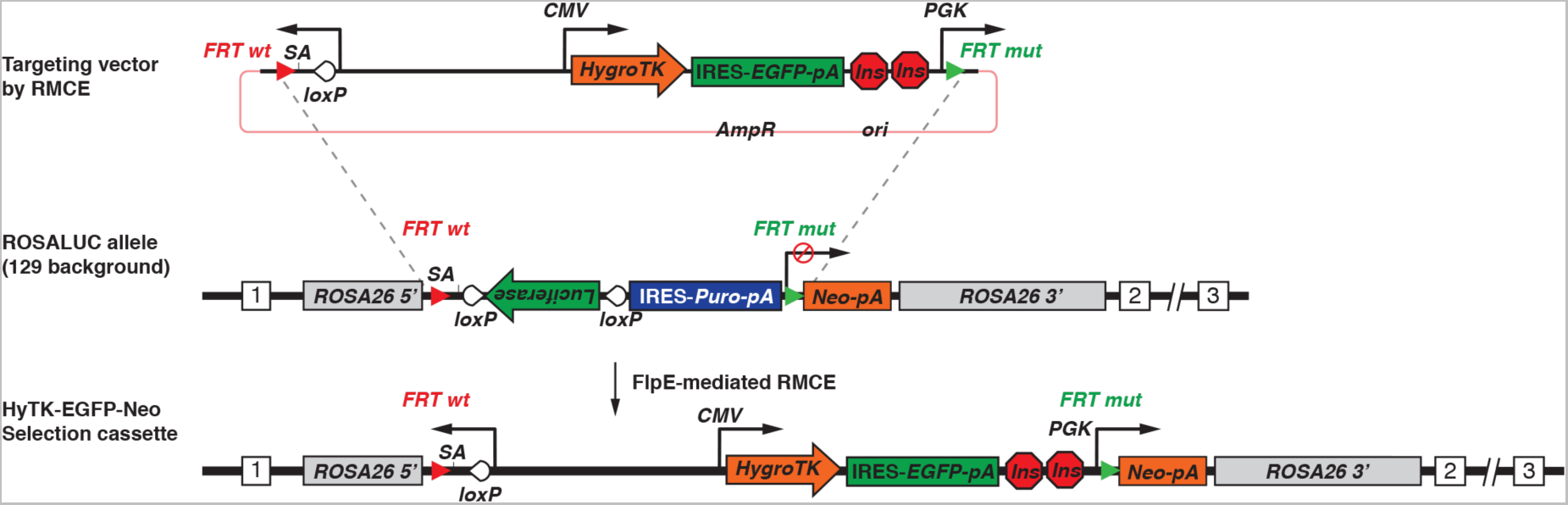
Site-specific integration of a versatile selection reporter cassette into the G4 ROSALUC ES cell line. Utilizing the recombination-mediated cassette exchange (RMCE) technique, the targeting vector was inserted by a *Flp*-recombinase into the ROSALUC allele as previously described (39). The vector introduced the hygromycin phosphotransferase-thymidine kinase (HyTK) fusion selectable marker, the enhanced green fluorescent protein (EGFP) and the phosphoglycerate kinase 1 (PGK) promoter, thus restoring the expression of the latent neomycin resistance gene upon the successful integration of the vector into the ROSALUC allele. Figure modified from (39).

**Fig. 1–Figure Supplement 2.**
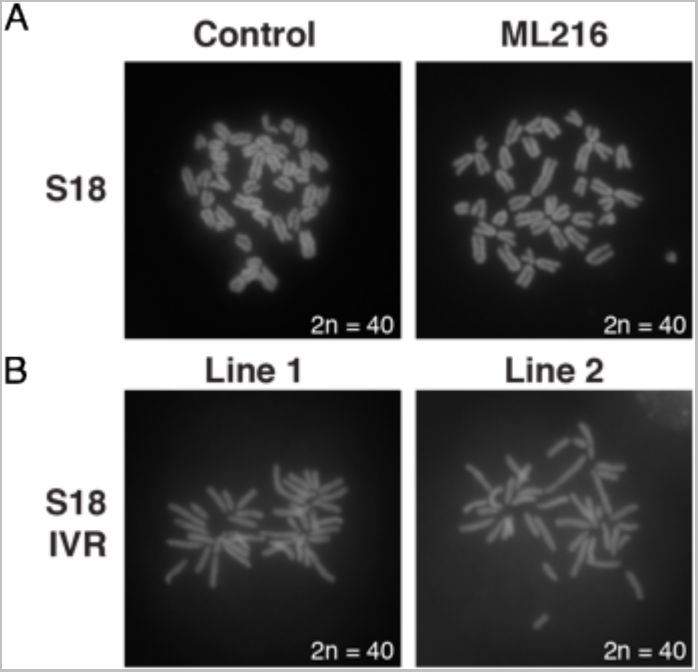
Normal karyotypes were maintained under culturing and IVR treatment. **(A)** Representative metaphase spreads from S18 line under control and ML216 treatment show normal karyotype of 2n = 40. **(B)** After confirmed IVR treatment, selected lines 1 and 2 were chosen for re-derivation. The karyotypes of both lines are also normal with high levels of euploidy. Whole embryos derived from laser-assisted morulae injection results showed that the S18 line, and IVR lines 1 and 2 are broadly competent to differentiate into diverse cell lineages (Fig. 4, Fig. 4–Figure Supplements 1 & 2).

**Fig. 2–Figure Supplement 1.**
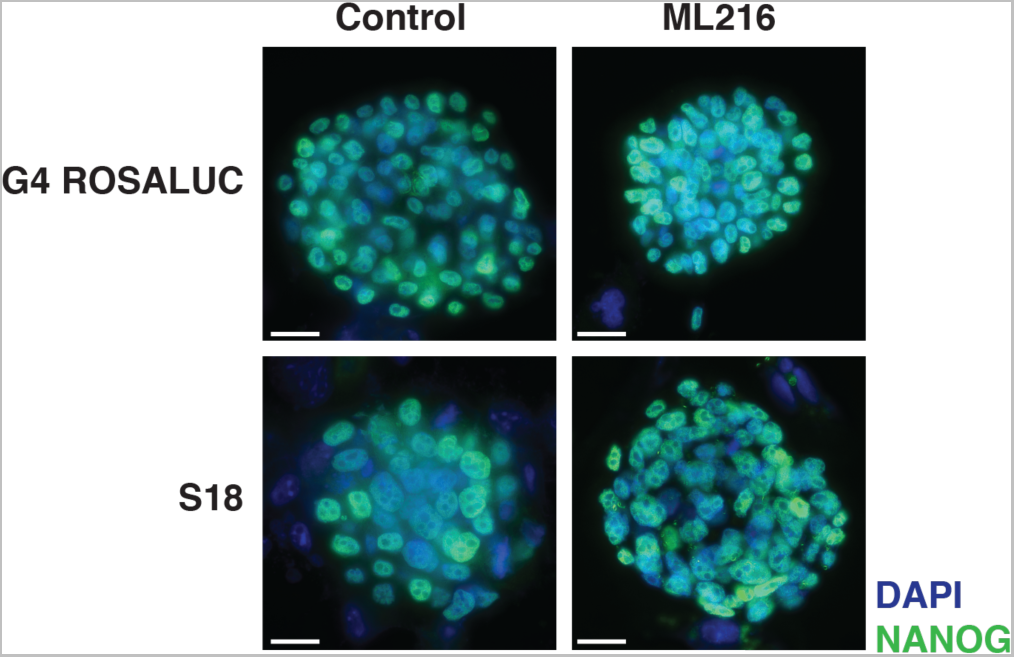
ML216 treatment is compatible with ES cell culturing. To determine if ML216 treatment affects ES cell colony viability and maintenance of stemness, we cultured ES cells strains G4 Rosaluc [(BL6 x 129S) F1] and S18 [(BL6 x *SPRET*) F1] under control 2i/LIF and 25 μM ML216 plus 2i/LIF conditions for 3 days. Both control and ML216 treated colonies showed good colony morphology, cell density and robust stem cell marker NANOG expression in both ES cell lines. We concluded that ML216 induction of *in vitro* recombination is compatible with ES cell culturing across considerable evolutionary divergence. Scale bar = 200 m.

**Fig. 2–Figure Supplement 2.**
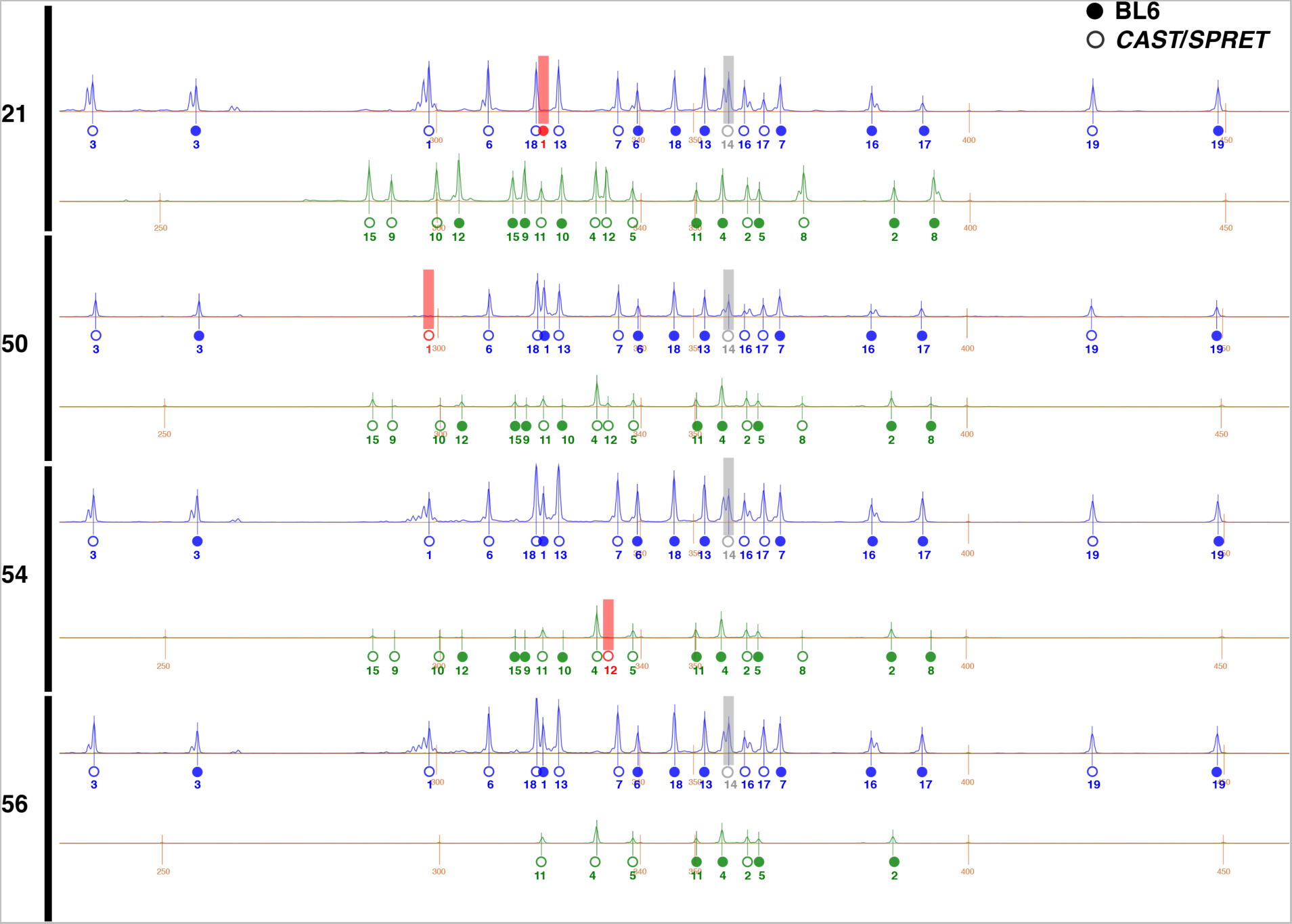
Multiplexed PCR genotyping screen for spontaneous recombinants. Hybrid ES cells [(BL6 x *CAST*) F1 hybrid line “E14”] were treated with ML216 and screened by multiplexed PCR genotyping at diagnostic markers within 10Mbp of each autosome chromosome (see Methods & Table S1). Amplified fragment sizes were determined using a capillary sequencer. The markers were designed such that they show staggered fragment sizes, allowing clear identification using fragment analysis software. Shown above are the electropherogram traces corresponding to the clones shown in Fig. 2, out of 46 total clones. The blue (FAM) and green (HEX) channels are shown separately for each sample, adjusted according to size standards (LIZ, orange, in basepairs. Fluorescence levels are shown on arbitrary units on the Y-axis). Genotype calls corresponding to BL6 (solid circles) and *CAST* or *SPRET* (open circles) alleles for each chromosome are shown underneath the called peaks (markers were designed for both outgroups. Only E14 analyses are included in this study). Missing genotypes indicative of recombination or LOH events are indicated in red. Chromosome 14 calls were removed due to invariant calls in all samples, including untreated F1 hybrid cells. This approach allowed us to rapidly screen through many colonies to detect possible recombinants. Notably, whole genome sequencing results suggested that in addition to the typical recombination events recovered by this multiplexed fragment analysis, further recombination events may occur elsewhere in the genome.

**Fig. 3–Figure Supplement 1.**
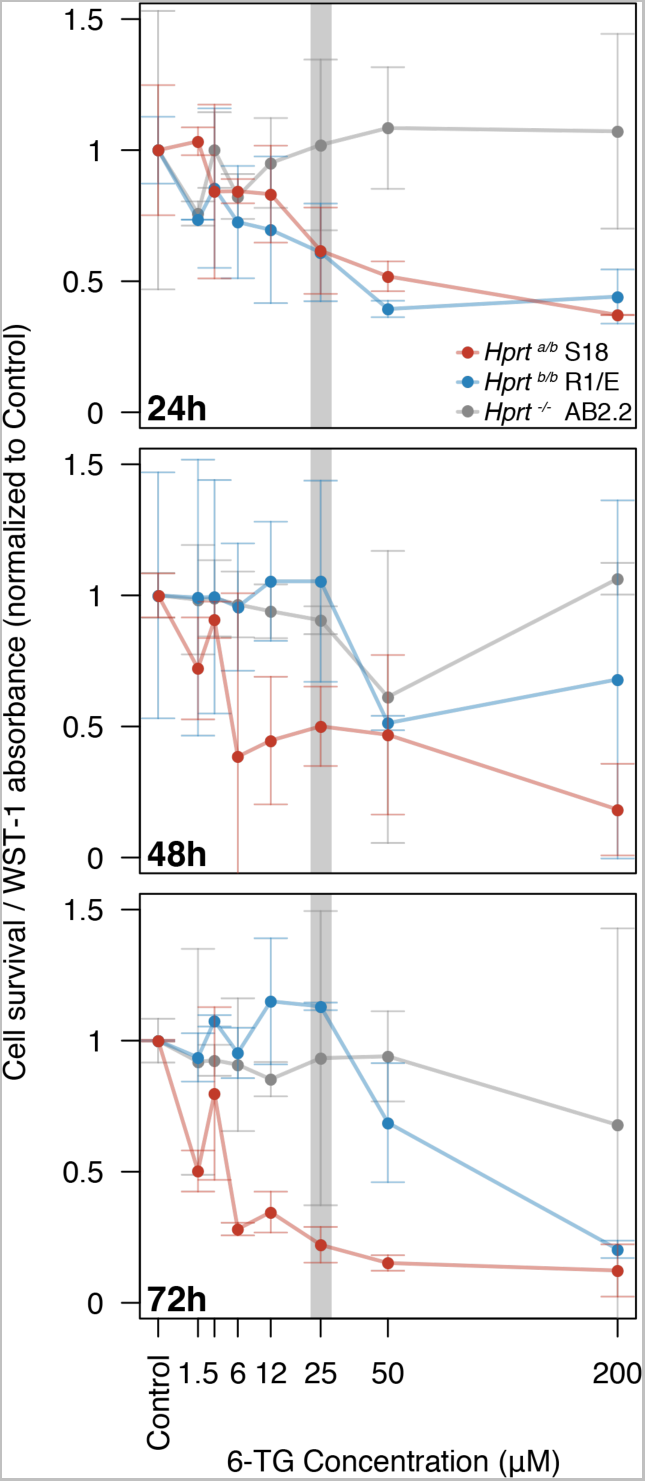
Optimal 6-TG concentration for differential *Hprt*-dependent cytotoxicity. Concentration for 6-TG treatment was determined by treating ES cells with concentrations ranging from 1.5 to 200 μM. Cell survival were determined by a colorimetric WST-1 absorbance assay. ES cells carrying different *Hprt-a*, *-b* or null alleles on various genetic backgrounds were assayed in duplicates over 24, 48 or 72 hours (*Hprt^a/b^* on (BL6 x *SPRET*) F1 S18 background: red; *Hprt^b/b^* on R1/E 129X1/129S1 background: blue; and *Hprt^-/-^* on AB2.2 129S5 background: grey). Absorbance values were normalized against control treatment of no 6-TG after subtracting blank measurements. We chose 25 NM 6-TG treatment for subsequent experiments for the strong survival difference between cells carrying *Hprt^a^* and those carrying *Hprt^b^* or null genotypes after 48 h. To ensure genome integrity for sequencing in flow mapping, we performed FACS already after 24 h of 25 μM 6-TG treatment together with a more sensitive DAPI exclusion cell viability assay. Plotted values are normalized mean between replicates ± s.d.

**Fig. 3–Figure Supplement 2.**
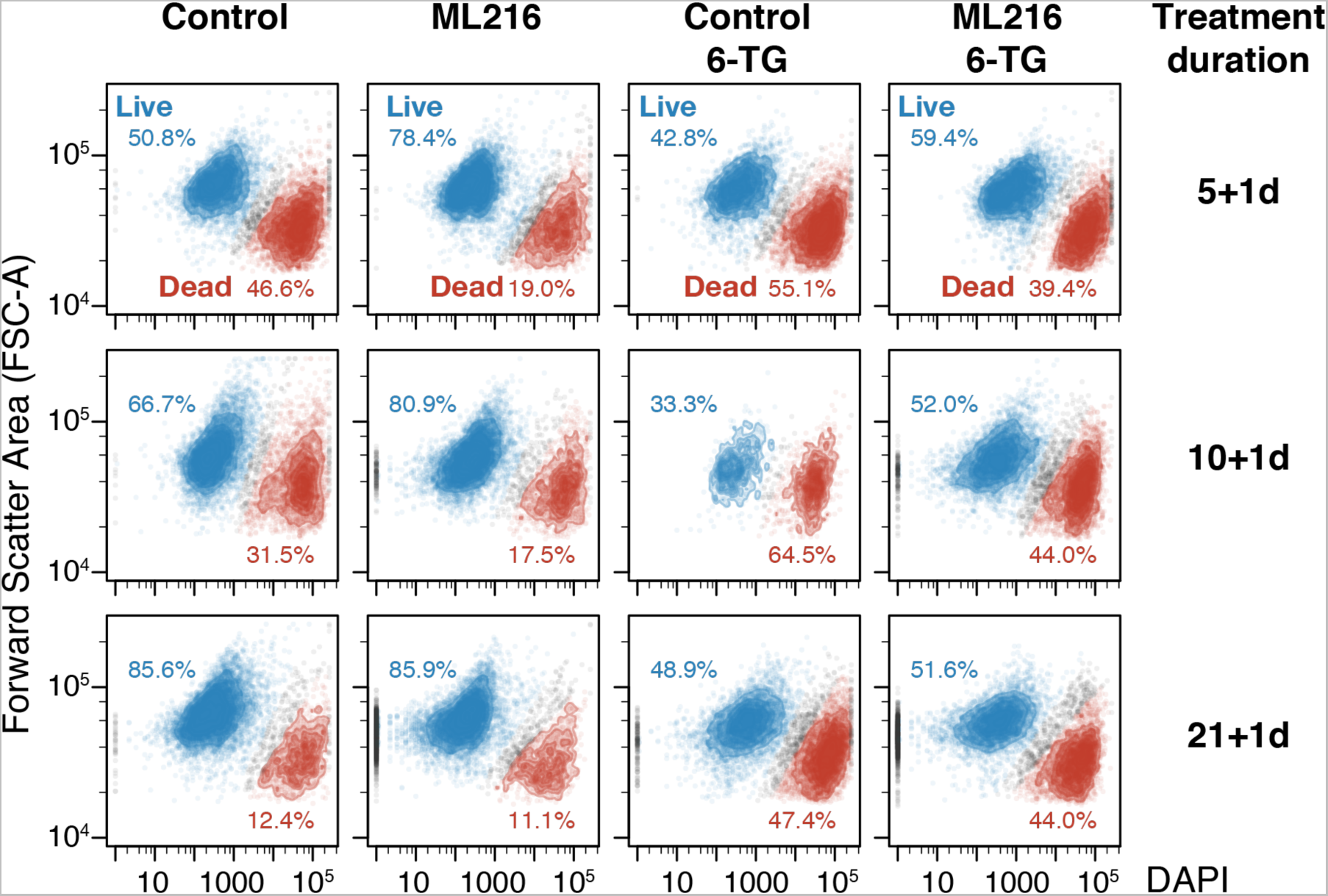
ML216 treatment maintains cell viability. S18 cells under various treatments were analyzed by flow cytometry to determine if ML216 (25 μM) treatment induces cell death. Under ML216 treatment alone, cells show robust viability (second column). Only after 1 d 6-TG treatment do the cells exhibit greatly increased cell death, as shown by the increased proportion of the “Dead” population (third column; red). Notably, combined ML216 and 6-TG treatment appears to mitigate cell damage and death, as indicated by the increased proportion of the “Live” population (third vs. fourth column; blue).

**Fig. 4–Figure Supplement 1.**
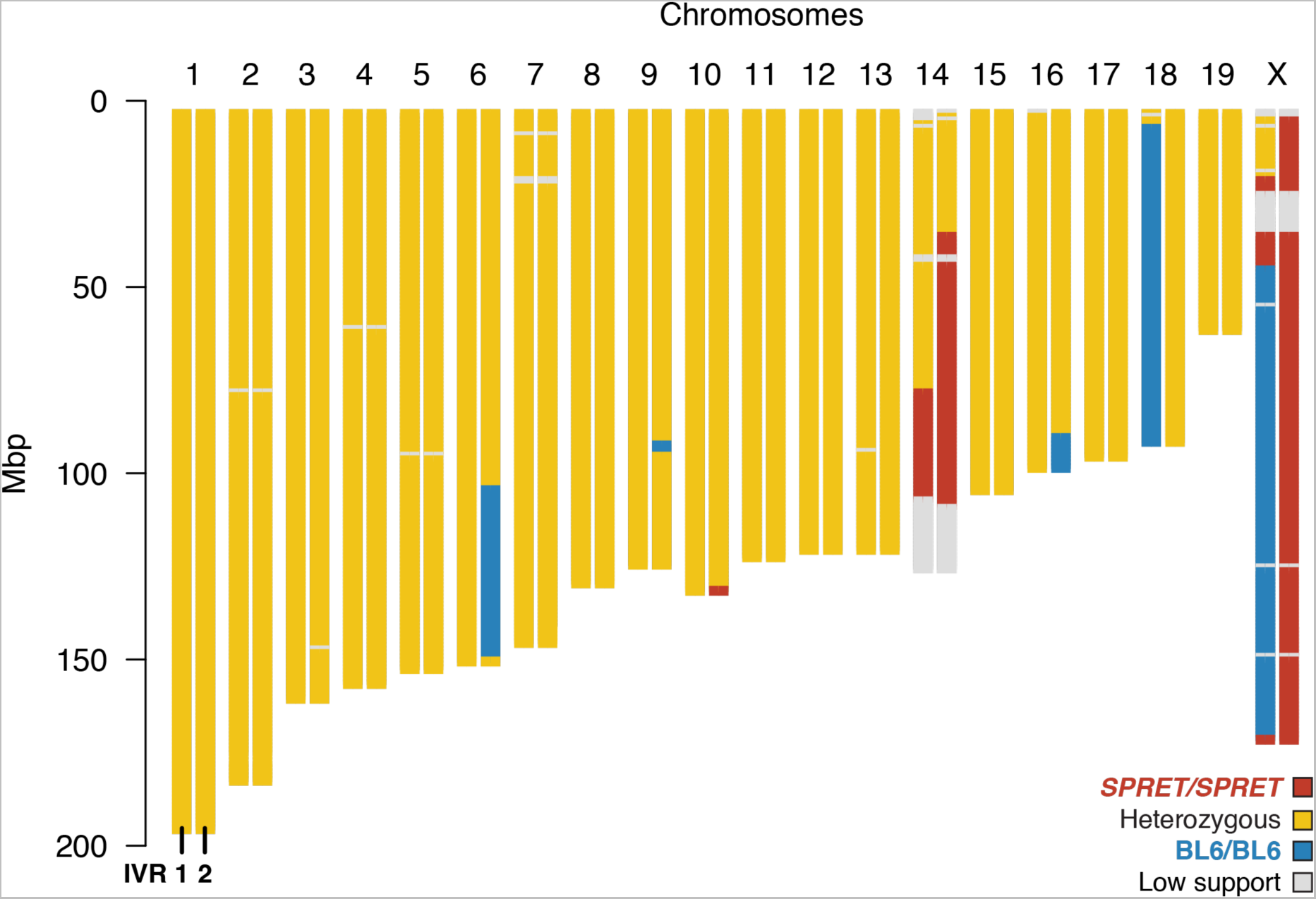
Genome-wide genotype of the two S18 IVR ES cell lines selected for embryo re-derivation. High-confidence genotypes of each line for each chromosome are plotted as heterozygous (yellow) and the two BL6/BL6 (blue) and *SPRET*/*SPRET* (red) homozygous genotypes. Low-coverage or repetitive regions were considered ambiguous (grey). Both lines 1 and 2 showed substantial proportion of the genome carrying heterozygous genotypes, reflecting their F1 hybrid origin. Because these lines were obtained through *6-TG* selection, much of the observed recombinant genotypes belong to Chromosome X. In addition, we have observed chromosome instability at the distal end of Chromosome 14 (also see Fig. 2–Figure Supplement 2). In addition, there are major genotypic differences between IVR lines 1 and 2 on chromosomes 6, 16 and 18, as well as X. Such recombinant genotype would be difficult, if not impossible to obtain under conventional breeding. These results illustrate the potential of applying IVR at expanded scale to investigate the genetic basis of species divergence.

**Fig. 4–Figure Supplement 2.**
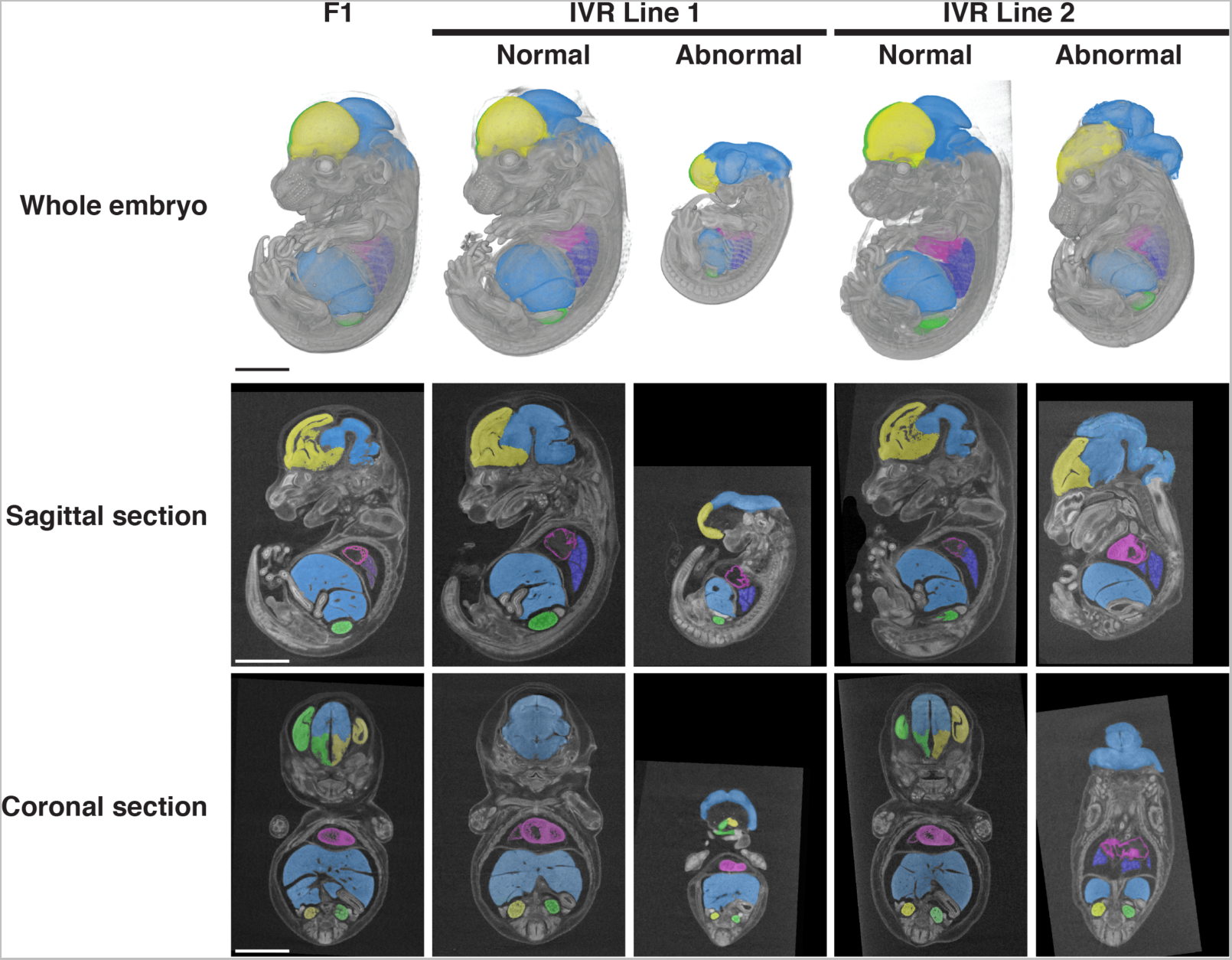
Whole embryos derived from F1 hybrid S18 non-recombinant and IVR ES cells. Embryos with almost exclusively ES cell contribution could be generated in the founder generation via laser-assisted morula injection. This allowed phenotyping of organismal traits by circumvention of hybrid sterility. Embryos were dissected in mid-gestation stage (approximately 14.5 days post-coitus, or embryonic E14.5), contrast-stained and scanned using X-ray micro-computer tomography (microCT) at 9.4 micron (μm) resolution. The use of contrast staining allowed identification and precise measurements of individual organs (colorized here for clarity). Embryos from non-recombinant S18 ES cells (left column) and two IVR ES cell lines were examined (columns 2–3 and 4–5 respectively). Representative individuals displaying normal and abnormal developmental phenotypes are shown as whole embryos with representative sagittal and coronal sections. In contrast to non-recombinant S18-derived embryos, multiple embryos from each IVR lines showed major craniofacial and neural tube closure defects. Despite a small sample size, such occurrence was highly atypical. Notably, defects in cell migration and cell–cell communication are consistent with hybrid incompatibilities. Following speciation, divergent genotype combinations carried by the same individuals have not been subjected to selection for compatible functions. Consequently, hybrid incompatibilities often result in developmental defects. Derivation of embryos from panels of IVR ES cell lines may allow genetic dissection of developmental variation arising from evolutionary divergences.

## Supplementary Files

**Table 1.**
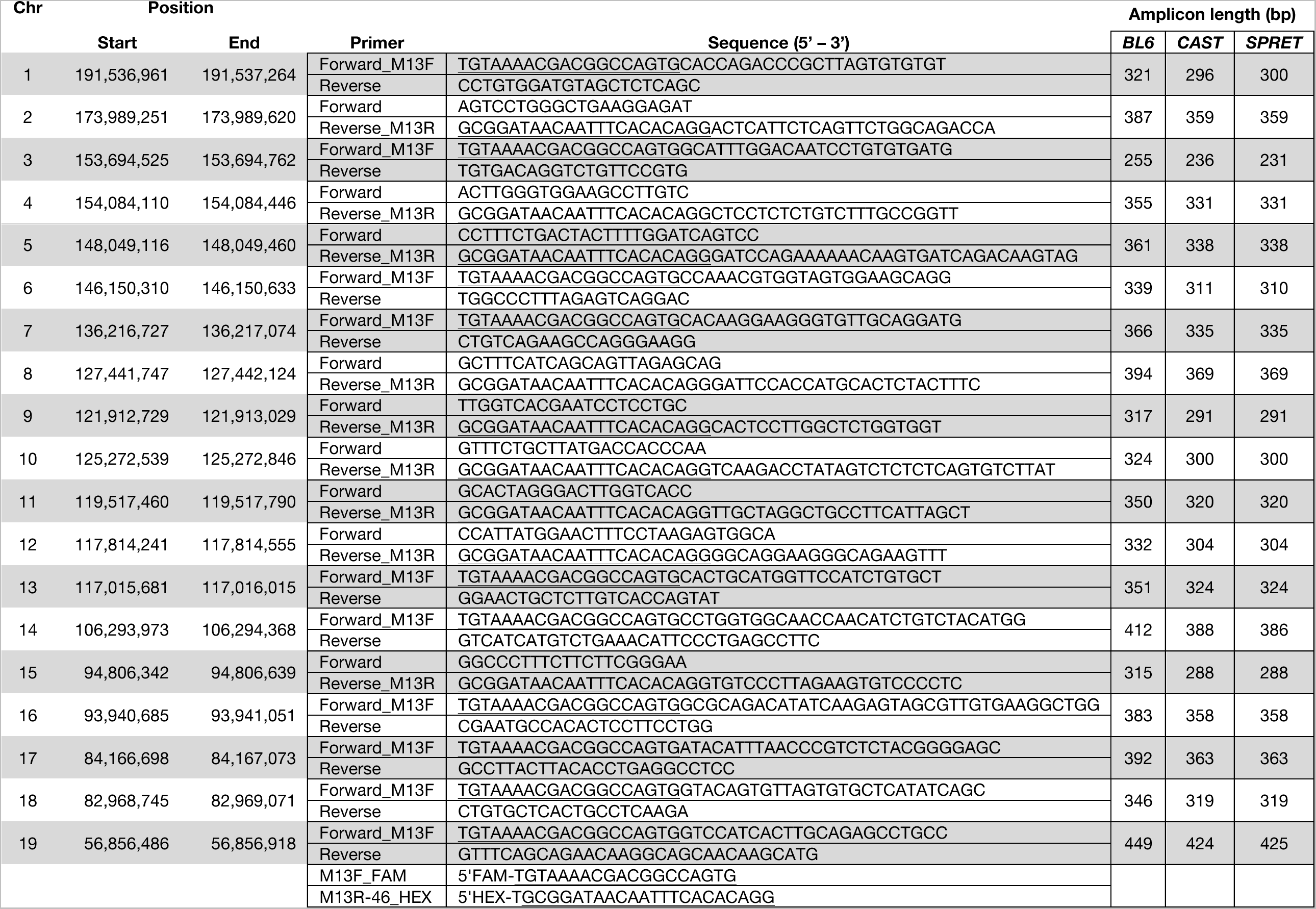
Oligonucleotide primers for multiplexed genotyping of sub-telomeric markers. Each pair of primers carry a M13-modified extension (underlined) to allow easy attachment of a third, universal fluorophore-conjugated primer for fragment analysis in a capillary sequencer as described in (43).

Multiplexed recombination detection PCR genotyping primers

## Supplementary Movies

Movie S1.

S18 (untreated) ES-cell derived embryo.

Whole embryos derived from F1 hybrid S18 non-recombinant ES cells. Embryos with almost exclusively ES cell contribution could be generated in the founder generation via laser-assisted morula injection. This allowed phenotyping of organismal traits by circumvention of hybrid sterility. Embryos were dissected in mid-gestation stage (approximately 14.5 d post-coitus, or embryonic E14.5), contrast-stained and scanned using X-ray micro-computer tomography (microCT) at 9.4 *μ*m resolution. The use of contrast staining allowed identification and precise measurements of individual organs (colorized here for clarity).

Movie S2.

S18 IVR Line 1 ES-cell derived embryos

Whole embryos derived from F1 hybrid S18 IVR Line 1 ES cells. Embryos with almost exclusively ES cell contribution could be generated in the founder generation via laser-assisted morula injection. This allowed phenotyping of organismal traits by circumvention of hybrid sterility. Embryos were dissected in mid-gestation stage (approximately 14.5 d post-coitus, or embryonic E14.5), contrast-stained and scanned using X-ray micro-computer tomography (microCT) at 9.4 *μ*m resolution. The use of contrast staining allowed identification and precise measurements of individual organs (colorized here for clarity). Representative individuals displaying normal and abnormal developmental phenotypes are shown.

Movie S3.

S18 IVR Line 2 ES-cell derived embryos

Whole embryos derived from F1 hybrid S18 IVR Line 2 ES cells. Embryos with almost exclusively ES cell contribution could be generated in the founder generation via laser-assisted morula injection. This allowed phenotyping of organismal traits by circumvention of hybrid sterility. Embryos were dissected in mid-gestation stage (approximately 14.5 d post-coitus, or embryonic E14.5), contrast-stained and scanned using X-ray micro-computer tomography (microCT) at 9.4 *μ*m resolution. The use of contrast staining allowed identification and precise measurements of individual organs (colorized here for clarity). Representative individuals displaying normal and abnormal developmental phenotypes are shown.

